# Combining landscape genomics and ecological modelling to investigate local adaptation of indigenous Ugandan cattle to East Coast fever

**DOI:** 10.1101/315184

**Authors:** Elia Vajana, Mario Barbato, Licia Colli, Marco Milanesi, Estelle Rochat, Enrico Fabrizi, Christopher Mukasa, Marcello Del Corvo, Charles Masembe, Vincent Muwanika, Fredrick Kabi, Tad Stewart Sonstegard, Heather Jay Huson, Riccardo Negrini, Stéphane Joost, Paolo Ajmone-Marsan, on behalf of The NextGen Consortium^^^

## Abstract

East Coast fever (ECF) is a fatal sickness affecting cattle populations of eastern, central, and southern Africa. The disease is transmitted by the tick *Rhipicephalus appendiculatus*, and caused by the protozoan *Theileria parva parva*, which invades host lymphocytes and promotes their clonal expansion. Importantly, indigenous cattle show tolerance to infection in ECF-endemically stable areas. Here, the putative genetic bases underlying ECF-tolerance were investigated using molecular data and epidemiological information from 823 indigenous cattle from Uganda. Vector distribution and host infection risk were estimated over the study area and subsequently tested as triggers of local adaptation by means of landscape genomics analysis. We identified 41 and seven candidate adaptive loci for tick resistance and infection tolerance, respectively. Among the genes associated with the candidate adaptive loci are *PRKG1* and *SLA*2. *PRKG1* was already described as associated with tick resistance in indigenous South African cattle, due to its role into inflammatory response. *SLA*2 is part of the regulatory pathways involved into lymphocytes’ proliferation. Additionally, local ancestry analysis suggested the zebuine origin of the genomic region candidate for tick resistance.

**Author summary:** The tick-borne parasite *Theileria parva parva* infects cattle populations of eastern, central and southern Africa, by causing a highly fatal pathology called “East Coast fever”. The disease is especially severe for the exotic breeds imported to Africa, as well as outside the endemic areas of East Africa. In these regions, indigenous cattle populations can survive to infection, and this tolerance might result from unique adaptations evolved to fight the disease. We investigated this hypothesis by using a method named “landscape genomics”, with which we compared the genetic characteristics of indigenous Ugandan cattle coming from areas at different infection risk, and located genomic sites potentially attributable to tolerance. In particular, the method pinpointed two genes, one (*PRKG1*) involved into inflammatory response and potentially affecting East Coast fever vector attachment, the other (*SLA2*) involved into lymphocytes proliferation, a process activated by *T. parva parva* infection. Our findings can orientate future research on the genetic basis of East Coast fever-tolerance, and derive from a general method that can be applied to investigate adaptation in analogous host-vector-parasite systems. Characterization of the genetic factors underlying East Coast-fever-tolerance represents an essential step towards enhancing sustainability and productivity of local agroecosystems.

## Introduction

East Coast fever (ECF) is an endemic vector-borne disease affecting cattle populations of eastern and central Africa. ECF etiological agent is the protozoan emo-parasite protozoan *Theileria parva* Theiler, 1904, vectored by the hard-bodied tick vector *Rhipicephalus appendiculatus* Neumann, 1901. The disease is reported to cause morbidity in indigenous populations and high mortality rates among exotic breeds and crossbreds, thus undermining the livestock sector development in the affected countries [1–3].

Cape buffalo (*Syncerus caffer* Sparrman, 1779) is *T. parva* native host, being its wild and asymptomatic reservoir [4]. A primordial contact between buffalo-derived *T. parva* and domestic bovines is likely to have occurred ~4.500 years before present (YBP) [5]. However, it is hard to define if the host jump affected taurine- or indicine-like cattle first, since no consensus can easily be reached to define who among *Bos taurus* and *B. indicus* migrated into ECF endemic regions first [6–8]. Indeed, African taurine cattle might have reached eastern Africa sometime between ~8,000 and 1,500 YBP [7,8], and the most ancient zebuine colonization wave is estimated to have occurred between ~4,000-2,000 YBP from the Asian continent, as suggested by the first certain archaeological record dated 1,750 YBP [6]. Once *T. parva* spread to domestic populations, coevolution between the parasite and the new hosts likely led to the divergence between buffalo- (*T. parva lawracei*) and cattle-specific (*T. parva parva*) parasite strains [9,10], and to the appearance of infection-tolerant indigenous herds [11,12].

Most likely, ECF-tolerance appeared (and is only observable) in areas where environmental conditions guaranteed a constant coexistence between the vector, the parasite and the domestic host. Such a particular situation, together with the evolution of some sort of “innate resistance” [13], plausibly prompted the establishment of an epidemiological state referred to as endemic stability, a condition where hosts become parasite reservoirs with negligible clinical symptoms [14]. However, no clear indication for a genetic control has been provided for ECF-tolerance so far [12], despite host genetic factors were identified for tolerance to tropical theileriosis [15] (a disease caused by the closely related *T. annulata*), and tick resistance [16].

Here, we propose an integrated approach based on ecological modelling and landscape genomics to explore the putative adaptive component sustaining ECF-endemic stability. Furthermore, given the existence of host populations showing differential susceptibility to ECF [13,14,17], we will refer to ECF-tolerance as a potential case of local adaptation [18]. Since endemically stable areas are currently inhabited by the indigenous zebu and the zebu *x* African *B. taurus* crosses sanga and zenga [8,19], two basic hypotheses can be associated to the origin of local adaptation to ECF: (i) at first adaptation appeared in local African *B. taurus* populations and was then introgressed into zebu and derived sanga and zenga crossbreds; alternatively, (ii) it appeared in *B. indicus*, and then either evolved independently in zebuine populations of eastern Africa, or was imported from the Indian continent, where similar selective pressures are recorded [20,21].

Specific regions in the South-West [14] and in the East [17] of current Uganda are reported to be ECF-endemically stable, thus making this country a candidate for investigating local adaptation to ECF. Moreover, indigenous Ugandan cattle populations are proven to be connected by high rates of gene flow [22], and a strong spatially varying selection is expected on their genomes because of regional climatic differences shaping ECF epidemiology over the country [11]. These requirements are all likely to have promoted local adaptation to the disease [23], even over short time scales (i.e. from thousands of years to few decades), as observed for other plant and animal species [24–26].

To test our approach, we exploited genomic data from indigenous Ugandan cattle and spatial information on parasite and vector occurrence. First, we modelled ECF-vector potential distribution and infection risk in cattle to define the spatially varying selective pressure over the host genomes. Then, we searched for Single Nucleotide Polymorphisms (SNPs) potentially involved into local adaptation to ECF through genotype-environment association (GEA) analysis, and we annotated candidate genes. Finally, we studied the ancestral origin of the identified genomic regions by means of local ancestry analysis to shed light on the possible evolutionary origins of local adaptation to ECF.

## Materials and Methods

### Ecological modelling

*R. appendiculatus* occurrence probability (*Ψ_R_*) and *T. parva parva* infection risk in cattle (*γ*) were modelled and used as environmental predictors into landscape genomics models. Geographical variability in both *Ψ_R_* and *γ* was assumed to describe the spatially heterogeneous selective pressure on cattle genomes. Further, *S. caffer* occurrence probability (*Ψ_S_*) was estimated and used in combination with *Ψ_R_* to model *γ*, as the geographical proximity between Cape buffaloes and cattle herds constitutes a factor for explaining ECF incidence. The following three sections will describe data and methods used to estimate *Ψ_R_*, *Ψ_S_*, and *γ*.

#### Raster data

Bioclimatic variables (BIO) referring to the time span between 1960 and 1990 were collected from the WorldClim database (v.1.4. release3) [27] at a spatial resolution of 30 arc-seconds and in the un-projected latitude/longitude coordinate reference system (WGS84 datum). Altitude information was collected from the SRTM 90m Digital Elevation Database (v.4.1) [28], which provides tiles covering Earth’s land surface in the WGS84 datum, at 90 m resolution at the equator. Altitude was used to compute terrain slope through the function `terrain` implemented in the R package `raster` [29]. The ten-year (2001-2010) averaged Normalized Difference Vegetation Index (NDVI) was derived for 72 ten-day annual periods from the “eMODIS products” (S1 Text) [30], in the WGS84 datum, and at a resolution of 250 m at the equator. A raster file describing cattle density (number of animals/km^2^) was acquired from the Livestock Geo-Wiki database [31], in the WGS84 datum, at a resolution of 1 km^2^ at the equator. A raster file describing each pixel distance from the nearest water source was obtained with the function `distance` within the R package `raster`. The “Land and Water Area” dataset from the Gridded Population of the World collection (GPV v.4) [32] was used to define water bodies in Uganda at a resolution of 30 arc-seconds with WGS84 datum.

All raster files were transposed into Africa Albers Equal Area Conic projection to guarantee a constant pixel size and meet the main assumption of the statistical technique used to model *Ψ_R_* and *Ψ_S_*, i.e. that each pixel presents the same probability to be randomly sampled in order to detect the species occurrence [33]. Raster files were standardised to the same resolution (~0.85 km^2^), origin, and extent. To avoid the inclusion of potentially misleading background locations while characterizing the occurrence probability of terrestrial species, inland water surfaces were masked prior to *Ψ_R_* and *Ψ_S_* estimation [34]. Quantum GIS (v.2.16.2) [35] and the R package `raster` were used for raster files manipulation.

#### Species distribution models

The R package `Maxlike` [36] was used to model *Ψ_R_* and *Ψ_S_* over Uganda. `Maxlike` is able to estimate species occurrence probability (*Ψ*) from presence-only data, by maximizing the likelihood of occurrences under the logit-linear model [33]:

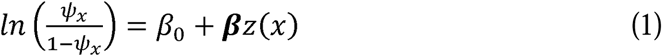

where *Ψ_x_* denotes the species occurrence probability in the *x* pixel of the landscape, *β_0_* the model intercept (i.e. the expected prevalence across the study area), ***β*** the vector of slope parameters, and *z(x*) the vector of environmental variables for *x*. Occurrence probability in *x* is derived from the inverse logit:

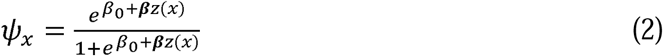

Fifty-one and 61 spatial records of *R. appendiculatus* and *S. caffer* (Figs 1A and 1B) were obtained from a tick occurrence database previously collected [37], and the Global Biodiversity Information Facility [38], respectively.

**Fig 1.**
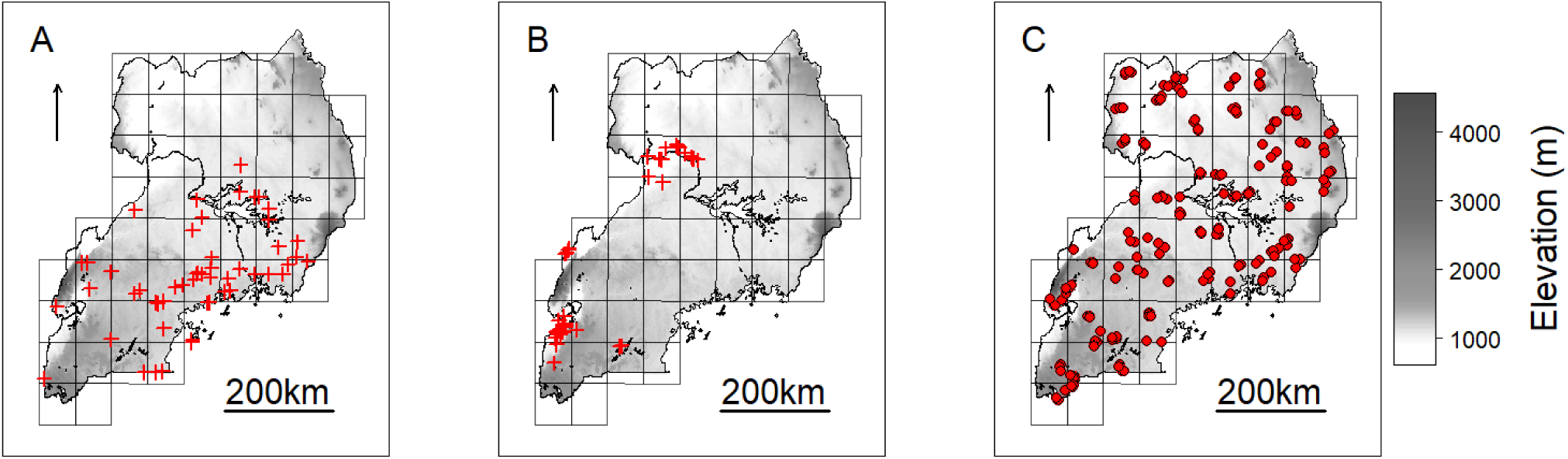
Occurrence records for species distribution modelling and NextGen sampling scheme. Spatial records (red crosses) used to estimate *R. appendiculatus* (A) and *S caffer* (B) distributions over Uganda, as derived from [37] and [38], respectively. Farms where cattle have been sampled to be genotyped and tested for *T. parva parva* infection are represented with red circles (C). The grid scheme used to sample farms during the NextGen project is shown on the background of each map (see main text), together with elevation.

The most relevant environmental variables affecting tick and Cape buffalo distributions were identified from the literature. Specifically, the BIO variables representing temperature/precipitation interaction in the most extreme periods of the year were used to model *R. appendiculatus* occurrence (Table 1 and S1 Fig.) [39,40], while altitude, terrain slope, NDVI, distance to water sources (Wd), and annual precipitation (BIO_12_) were used to model the Cape buffalo distribution [41–43]. A `Maxlike` regression analysis was applied to individuate the NDVI values best predicting the available *S. caffer* occurrences, and the period April 6-15 was retained for subsequent analyses (S2 Fig.). No variable depicting the top-down regulatory effect of predators on buffalo populations was considered, as bottom-up ecological mechanisms (like quantity and quality of food resources) are argued to play the main role in determining large herbivores distribution [44].

**Table 1.**
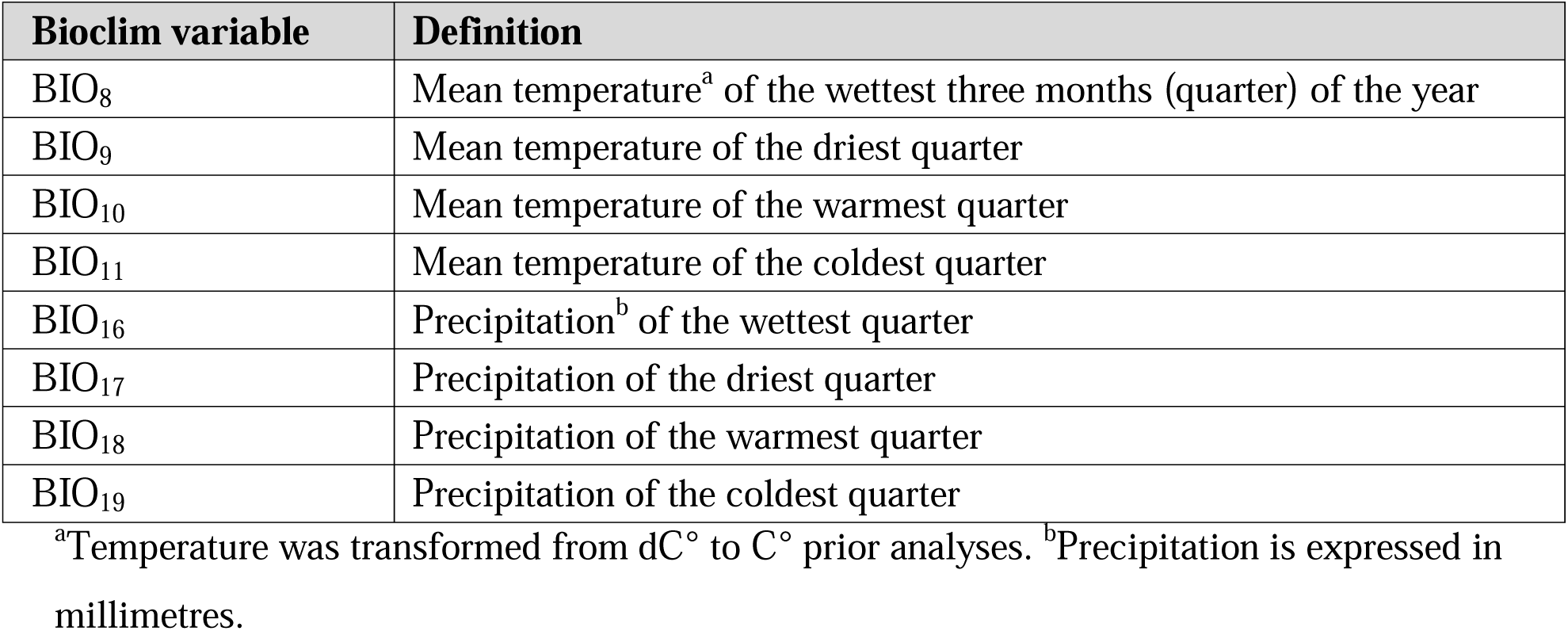
Predictors for *R. appendiculatus* distribution model.

Collinearity was checked prior to analyses by computing pairwise absolute correlations (|r|) between variables, which were considered collinear when |r| exceeded the suggested threshold of 0.7 [45]. High collinearity was found among BIO variables, which were then subjected to principal components analysis (PCA) to obtain orthogonal predictors for *Ψ_R_*.

Obtained components were tested into univariate and multivariate *R. appendiculatus* distribution models. Particularly, components explaining up to 95% of the original variance [46] were individuated and tested with different combinations into multivariate models, leading to a total of twelve candidate *R. appendiculatus* distribution models. Conversely, all the combinations of environmental variables were tested into univariate up to penta-variate Cape buffalo distribution models, resulting in a total of 31 candidate models for predicting *S. caffer* potential distribution.

In both cases, Bayesian Information Criterion (BIC) was used to select the best models [47]. Bring’s standardization [48,49] was applied on predictors before parameters’ estimate, and the delta method was implemented to compute the 95% confidence intervals around the fitted *Ψ_Rx_* and *Ψ_Sx_*.

#### Infection risk model

In the context of the European Project NextGen (http://nextgen.epfl.ch), 587 blood samples from Ugandan indigenous cattle were tested for the presence/absence of *T. parva parva* p104 antigen DNA sequence [11]. Samples were collected and georeferenced in correspondence of 203 farms distributed over a grid of 51 cells covering the whole Uganda, with an average of 12 (±4 s.d.) animals/cell, and three (±1 s.d.) animals/farm (Fig 1C).

ECF epidemiology is complex and determined by both biotic and abiotic factors [2]. Particularly, *R. appendiculatus* occurrence (*Ψ_R_*) [1, 50–52], cattle density (Cd) [3,53], potential proximity with *S. caffer* (*Ψ_S_*) and the maximal temperature in the warmest month of the year (BIO_5_) were considered to predict *γ*. BIO_5_ was used to account for the possible limiting effect of high temperatures on the parasite development into the tick [54]. Predictors’ values were obtained at the geographical position of each animal (i.e. locations of the farms), checked for the presence of collinearity (as done for the species distribution models) and outliers (S3 Fig.), and subsequently standardized following Bring’s procedure prior to parameters’ estimation.

Infection risk for any *i-th* animal was modelled using a binary mixed-effects logistic regression, where *Ψ_R_*, BIO_5_, Cd, and *Ψ_R_* were specified as fixed effects, and random intercepts were estimated for each farm to account for the possible influence of local environmental conditions and management practices (e.g. differential use of acaricides), as well as unmeasured biological features (e.g. breed- or individual-specific response to tick burden) [13]. Since geographical position of samples was recorded at the farm-level, all the animals coming from a given farm were characterized by equal environmental values. Thus, the model can be written as:

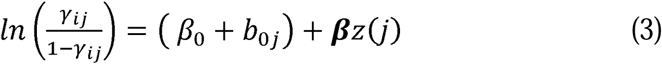

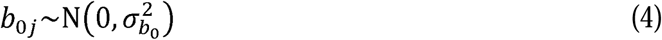

where *γ_ij_* represents *T. parva parva* infection risk for the *i-th* animal in the *j-th* farm, *β_0_* is the population intercept [55], *β_0_*+*b_0j_* is the *j-th* farm random intercept, ***β*** the vector of slope parameters, and *z*(*j*) the vector containing the predictors’ values as derived from the pixel where the *j-th* farm is located, equal for all the animals in *j*. In this way, animals in *j* are expected with the same predicted *γ*, so that infection risk in the *j-th* farm can be calculated using the population model from the previous equation:

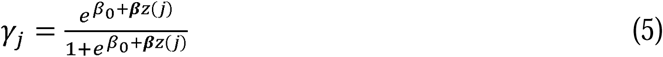

Estimates of the parameters were obtained through the Maximum Likelihood criterion using the glmer function included in the R package lme4 [56].

### Landscape genomics

#### Molecular datasets

The NextGen project genotyped 813 georeferenced indigenous cattle from Uganda using the medium-density BovineSNP50 BeadChip (54,596 SNPs; Illumina Inc., San Diego, CA, USA). Landscape genomics analyses were carried out on this set of animals, which will be referred to as the “landscape genomics dataset” (LGD). Samples were collected according to the spatial scheme represented in Fig 1C, and encompassed 503 of the individuals tested for *T. parva parva* infection. Quality control (QC) procedures were carried out with the software PLINK v.1.7 [57]. LGD was limited to autosomal chromosomes and pruned for minor allele frequency (MAF) <0.01, genotype call rates <0.95, and individual call rate <0.9. Pairwise genome-wide identity-by-descent (IBD) values were estimated, and one individual per pair showing IBD>0.5 was excluded from analyses to reduce the risk of spurious associations due to unreported kinship [58]. To avoid excluding too many individuals from nearby areas, spatial positions of the highlighted pairs were considered prior to removal.

Population genetic structure of Ugandan cattle was studied on the landscape genomics dataset merged with molecular data from other European taurine, African taurine, zebuine and sanga populations retrieved from various sources and for different geographical areas worldwide (S1 Table). This extended dataset will be referred to as the “population structure dataset” (PSD). PLINK was used to prune PSD for linkage disequilibrium (LD) >0.1 with sliding windows of 50 SNPs and step size of 10 SNPs (option`--indep-pairwise 50 10 0.1`), and to filter for the QC thresholds previously reported.

#### Population structure analysis

PSD was analysed with ADMIXTURE v.1.3.0 [59] for a dual purpose. Firstly, to provide genotype-environment association tests with population structure predictors in order to reduce the risk of false positive detections [60,61]. To this aim, we decided to use membership coefficients for the four-cluster solution (*K*=4), as this was reported to be the best partition based on the ADMIXTURE cross-validation index for the same set of Ugandan individuals undergoing landscape genomics in the present study [22]. Due to strong collinearity (|r|>0.7) [45] among the membership coefficients of two ancestral components, a PCA was performed trough the R function prcomp to obtain synthetic and orthogonal population structure predictors. Secondly, to identify the main gene pools present in Uganda in the context of a worldwide-extended dataset, and therefore guide selection of proper reference populations for local ancestry analysis.

#### Genotype-environment associations

We used the software SAMβADA v.0.5.3 [22,62] to test for associations between cattle genotypes and *Ψ_R_* and *γ* at sampling locations. Given diploid species and biallelic markers, SAMβADA runs three models per locus, one for each possible genotype.

Each model estimates the probability *π_i_* for the *i-th* individual to carry a given genotype, as a function of the considered environmental and population structure variables:

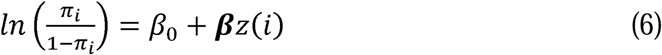

and thus:

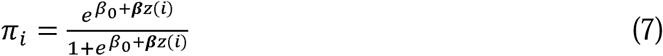

Genotype-environment association tests were carried out through a likelihood-ratio test comparing a null and an alternative model for each genotype [22]. Particularly, null models comprised the population structure predictors alone, while alternative ones included population structure predictors plus either *Ψ_R_* or *γ*. A genotype was considered significantly associated with *Ψ_R_* and/or *γ* if the resulting *p-value* associated with the likelihood-ratio test statistic (*D*) was lower than the nominal significance threshold of 0.05 after Benjamini-Hochberg (BH) correction for multiple testing (*H_0_*: *D*=0, *α*_BH_=0.05; S2 and S3 Text). The R function p.adjust was used to perform *p-values* corrections, and predictors were centred prior to analysis to ease estimation of model parameters.

#### Gene annotation

Global linkage disequilibrium (LD) decay was estimated using SNeP v.1.11 [63] to define LD extent around marker loci. A window of ±25 kbp (r^2^≈0.2) was then selected around those SNPs associated with *Ψ_R_* and/or *γ* to annotate genes in the Ensembl database release 87 [64]. Annotated genes were investigated for known biological function according to the literature, and candidate genes identified based on their pertinence with ECF local adaptation.

### Local ancestry

#### Molecular dataset

Target population for local ancestry analysis comprised 102 indigenous Ugandan cattle individuals collected during the NextGen sampling campaign (two animals sampled per cell; Fig 1C), and genotyped with the BovineHD BeadChip (777,961 SNPs; Illumina Inc., San Diego, CA, USA). Reference populations (see Results section) were selected in coherence with the major Ugandan gene pools identified by the ADMIXTURE analysis (S4 Text). Target and reference populations were pooled in a “local ancestry dataset” (LAD). Only autosomal SNPs passing the same filtering parameters applied to LGD were retained for analysis.

#### PCADMIX analysis

Local ancestry investigation allows to assign the ancestral origin of a chromosomal region (window) given two or more reference populations, and have been used to infer the admixture history of closely related groups [65], identify signals of adaptive introgression [66], and highlight target regions of recent selection [67]. Here, PCADMIX v.1.0 [68] was used to infer local genomic ancestry of the Ugandan samples. Given the SNPs density present in LAD (i.e. one SNP every ~3.4 kbp, on average), we used 20 SNPs per window to obtain a window size comparable to the optimal one suggested in [68].

#### Beta regression analysis

Genomic windows hosting SNPs in linkage with the candidate genes for local adaptation were identified and their ancestry proportions computed per sampling cell (Fig 1C). Average *Ψ_R_* and *γ* per cell values (hereafter *Ψ_Rc_* and *γ_c_*, respectively) were derived using the zonal.stats function included in the R package spatialEco [69]. In order to test for significant associations between ancestry proportions and *Ψ_Rc_* and *γ_c_*, a beta regression analysis was performed using the R package betareg [70], according to the model:

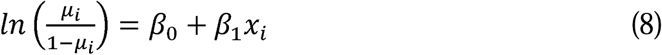

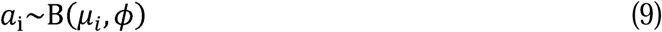

Where *a_i_* is the ancestry proportion observed in cell *i*, which is assumed to derive from a beta distribution B(μ_*i*_, □) with mean μ_*i*_=E(*a_i_*) and precision parameter □, *x_i_* is either average *Ψ_R_* or *γ* in cell *i*, *β_0_* and *β_1_* are intercept and regression coefficient, respectively. Expected ancestry proportion in *i* was calculated through the inverse logit:

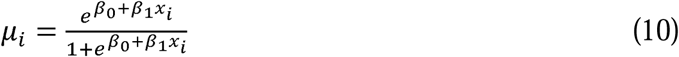

Ancestry proportions were transformed prior to analysis [71], and the Maximum Likelihood criterion was used to estimate model parameters.

### Ethics Statement

The NextGen sampling campaign was carried out during years 2011 and 2012, before Directive 2010/63/EU came into force (i.e., 1 January 2013). Thus, all experimental procedures were compliant with the former EU Directive 86/609/EEC, according to which no approval from dedicated animal welfare/ethics committee was needed for this study. The permission to carry out the study was obtained from the Uganda National Council for Science and Technology (UNCST) reference number NS 325 [11]. The permission to carry out the sampling at each farm was obtained directly from the owners.

## Results

### Ecological modelling

#### Species distribution models

The first three principal components (PC_1_, PC_2_, and PC_3_) accounted for more than 95% of the variance among the BIO predictors, and were subsequently tested into multivariate `Maxlike` models to estimate *Ψ_R_*. Particularly, PC_1_ (61%) was mainly correlated with BIO variables linked to temperature (BIO_8_, BIO_9_, BIO_10_ and BIO_11_), PC_2_ (19%) with precipitation (BIO_16_, BIO_17_, BIO_18_ and BIO_19_), and PC3 (15%) with both temperature and precipitation (BIO_19_ and BIO_8_) (Table 1 and S5 Fig.). The model employing PC_1_, PC_2_, and PC_3_ was selected based on the BIC metric (S6 Fig.), with PC_1_ and PC_2_ showing a significant positive effect on the tick distribution, and PC_3_ a significant negative effect (Table 2) (*H_0_*: *β_i_*=0, *α*=0.05). The model predicts low habitat suitability in the regions North of the Lakes Kwania, Kyoga and Kojwere (0<*Ψ_R_*<0.1), and favourable ecological conditions around Lake Victoria (0.4<*Ψ_R_*<1) and South-West of Lake Albert (0.4<*Ψ_R_*<0.8), these latter separated by a corridor of lower suitability (0<*Ψ_R_*<0.3) (Fig 2A and S7 Fig.).

**Fig 2.**
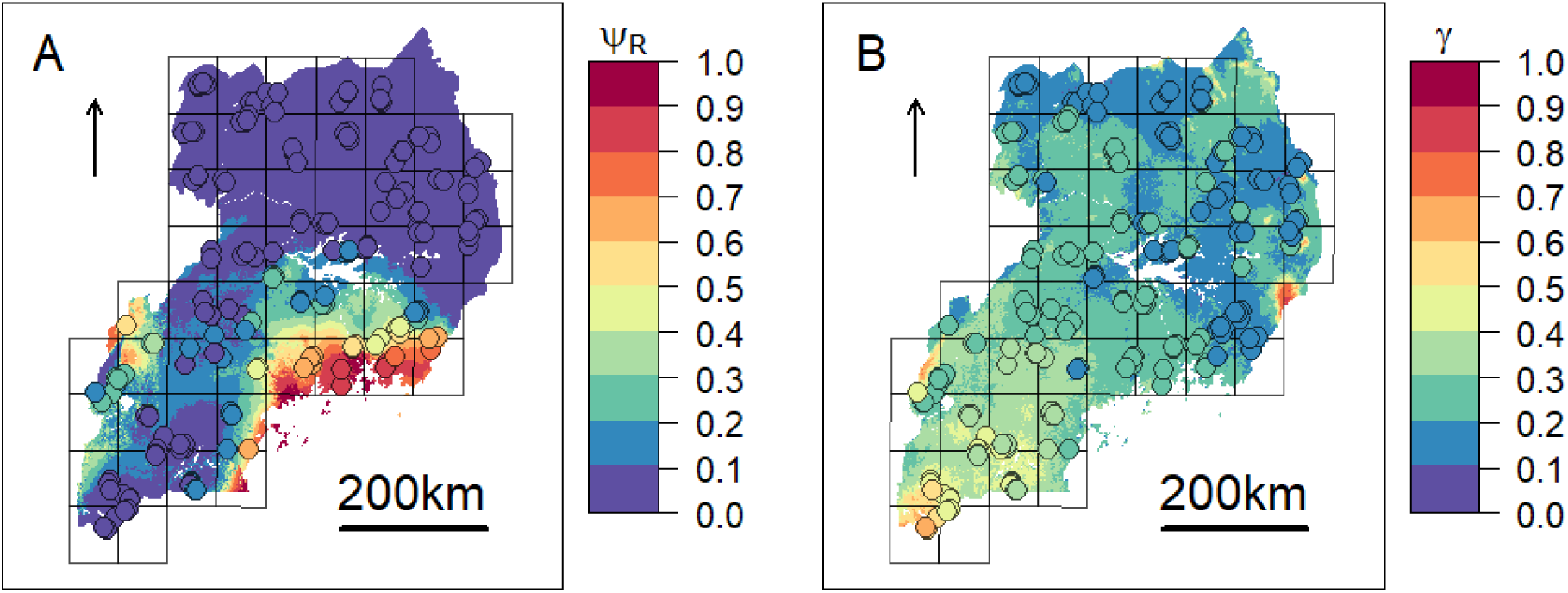
Predicted distributions ECF vector and infection risk. (A) *R. appendiculatus* occurrence probability (*Ψ_R_*) as predicted by the selected distribution model. (B) Predicted *T. parva parva* infection risk (*γ*). Colour from blue to red tones corresponds to increasing values of *Ψ_R_* and *γ*. Sampled farms are represented with circles, and coloured according to *Ψ_R_* and *γ* values estimated at their geographical location.

**Table 2.** Maxlike results for *R. appendiculatus* distribution model.

| Coefficient | Estimate | SE | *p*-value | OR<sup>a</sup> | OR<sub>low</sub><sup>b</sup> | OR<sub>up</sub><sup>c</sup> |
| --- | --- | --- | --- | --- | --- | --- |
| $\beta_0$ | -2.905 | 0.561 | 2.24E-07*** | 0.055 | 0.018 | 0.164 |
| PC<sub>1</sub> | 0.796 | 0.303 | 8.56E-03** | 2.217 | 1.224 | 4.014 |
| PC<sub>2</sub> | 0.822 | 0.37 | 2.62E-02* | 2.275 | 1.102 | 4.698 |
| PC<sub>3</sub> | -1.799 | 0.629 | 4.27E-03** | 0.165 | 0.048 | 0.568 |

Point estimates (Estimate) of the standardized regression coefficients (Coefficient) are reported on the logit scale together with their standard errors (SE), *p*-values and the associated odds ratios (OR). A significant effect is reported with \*\*\* when the *p*-value (*p*) associated to a regression coefficient is $\leq 0.001$ ; \*\* when $0.001 < p < 0.01$ ; \* when $0.01 < p < 0.05$ ; . when $0.05 < p < 0.1$ .
<sup>a</sup>Odds ratios associated to regression coefficients express the expected change in the ratio $\Psi_R/(1-\Psi_R)$ , for a one standard deviation increase of the concerned predictor, holding all the other predictors fixed at a constant value. <sup>b</sup>Odds ratio 95% confidence interval (CI), lower bound. <sup>c</sup>Odds ratio 95% CI, upper bound.

No excessive collinearity was recorded among the predictors for *Ψ_S_*. The best model according to the BIC metric included: altitude, annual precipitation, average NDVI and distance from the nearest water source (Table 3 and S8 Fig.). The model predicts the highest habitat suitability (0.2<*Ψ_S_*<0.8) in the near proximity of the water bodies (especially along the White Nile in the North-West, the south-eastern coasts of Lake Édouard, and the northern coasts of Lake George), and in small areas near the Katonga Game Reserve (S9 Fig.).

**Table 3.** Maxlike results for *S. caffer* distribution model.

| Coefficient | Estimate | SE | *p*-value | OR<sup>a</sup> | OR<sub>low</sub><sup>b</sup> | OR<sub>up</sub><sup>c</sup> |
| --- | --- | --- | --- | --- | --- | --- |
| $\beta_0$ | -9.130 | 0.790 | 6.46E-31*** | 0.000 | 0.000 | 0.001 |
| Altitude | −1.095 | 0.293 | 1.90E−04*** | 0.335 | 0.188 | 0.594 |
| BIO<sub>12</sub> | −0.800 | 0.180 | 9.03E−06*** | 0.449 | 0.316 | 0.639 |
| NDVI | 2.862 | 0.329 | 3.38E−18*** | 17.499 | 9.181 | 33.343 |
| Wd | −1.996 | 0.434 | 4.23E−06*** | 0.136 | 0.058 | 0.318 |

Point estimates (Estimate) of the standardized regression coefficients (Coefficient) are reported on the logit scale together with their standard errors (SE), *p-values* and the associated odds ratios (OR). Significant regression coefficients are highlighted with \*\*\* when their *p-values* (*p*) are $\leq 0.001$ ; \*\* when $0.001 < p \leq 0.01$ ; \* when $0.01 < p \leq 0.05$ ; . when $0.05 < p \leq 0.1$ .
<sup>a</sup>Odds ratios associated to regression coefficients express the expected change in the ratio $\Psi_S/(1-\Psi_S)$ , for a one standard deviation increase of the concerned predictor, holding all the other predictors fixed at a constant value. <sup>b</sup>Odds ratio 95% confidence interval (CI), lower bound. <sup>c</sup>Odds ratio 95% CI, upper bound.

#### Infection risk model

Following outliers inspection, *Ψ_R_*, Cd and *Ψ_S_* were transformed on the log_10_ scale to reduce the observed skewness in the distributions (S3 Fig.). No excessive collinearity was observed among the model predictors (|r|<0.7). All the explanatory variables except for Cd showed a significant effect (*H_0_*: *β_i_*=0, *α*=0.05) on infection risk. Particularly, BIO_5_ and Ψ*_R_* showed a negative association with *γ*, while *Ψ_S_* resulted positively associated (Table 4). Overall, northern regions of Uganda present a low probability of infection (0.1<*γ*<0.3). A similar range is observed southwards, in the region comprised between Lake Kyoga, Lake Victoria, Lake Albert and the eastern borders with Kenya. South-westwards, infection probability increases following a positive gradient from *γ* ≈ 0.30 to *γ* ≈ 0.70 in the most southern districts (Fig 2B).

**Table 4.**
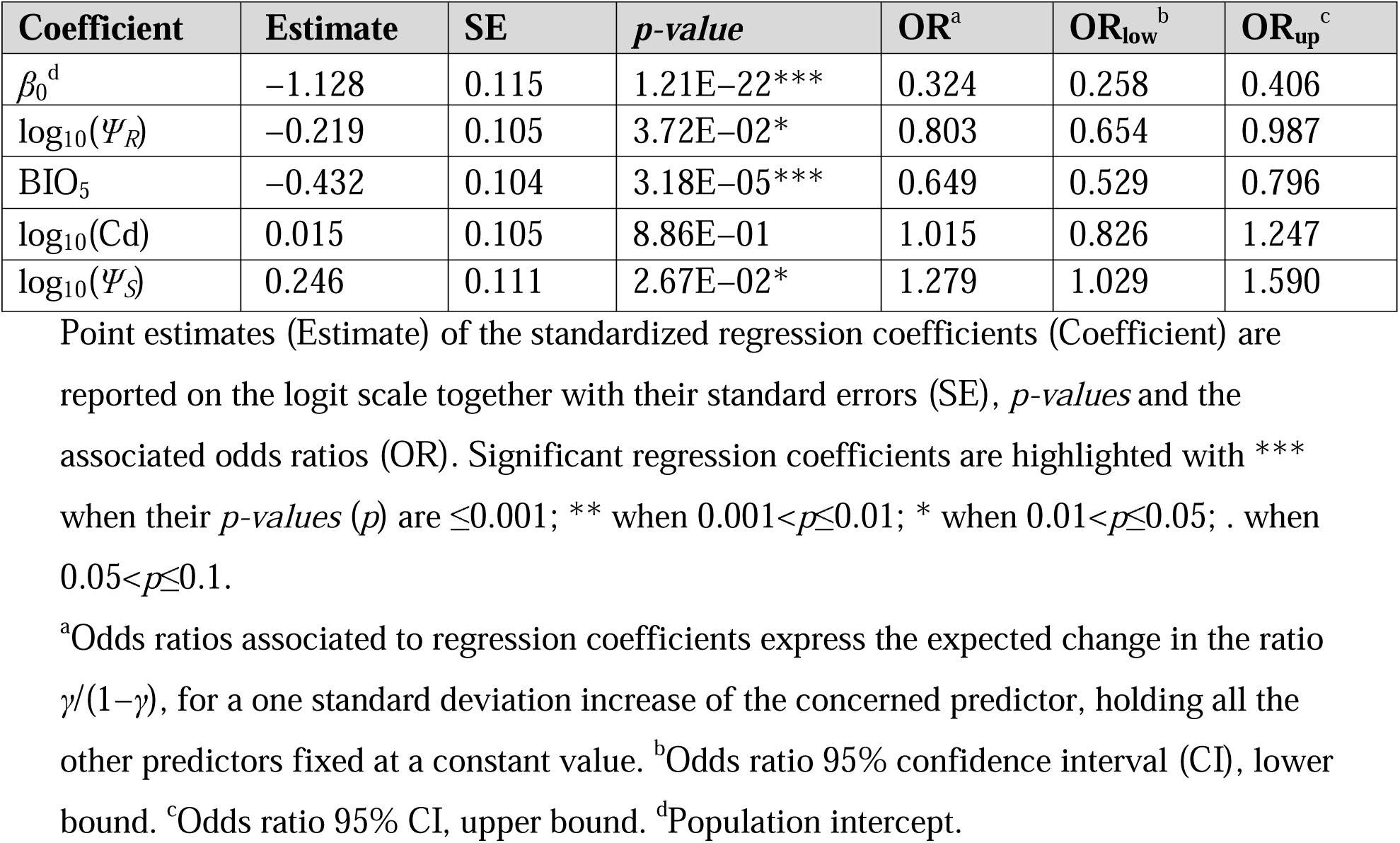
*T. parva parva* infection risk model results.

| Coefficient | Estimate | SE | *p*-value | OR<sup>a</sup> | OR<sub>low</sub><sup>b</sup> | OR<sub>up</sub><sup>c</sup> |
| --- | --- | --- | --- | --- | --- | --- |
| $\beta_0^d$ | -1.128 | 0.115 | 1.21E-22*** | 0.324 | 0.258 | 0.406 |
| $\log_{10}(\Psi_R)$ | -0.219 | 0.105 | 3.72E-02* | 0.803 | 0.654 | 0.987 |
| BIO<sub>5</sub> | -0.432 | 0.104 | 3.18E-05*** | 0.649 | 0.529 | 0.796 |
| $\log_{10}(\text{Cd})$ | 0.015 | 0.105 | 8.86E-01 | 1.015 | 0.826 | 1.247 |
| $\log_{10}(\Psi_S)$ | 0.246 | 0.111 | 2.67E-02* | 1.279 | 1.029 | 1.590 |

Point estimates (Estimate) of the standardized regression coefficients (Coefficient) are reported on the logit scale together with their standard errors (SE), *p*-values and the associated odds ratios (OR). Significant regression coefficients are highlighted with \*\*\* when their *p*-values (*p*) are $\leq 0.001$ ; \*\* when $0.001 < p \leq 0.01$ ; \* when $0.01 < p \leq 0.05$ ; . when $0.05 < p \leq 0.1$ .
<sup>a</sup>Odds ratios associated to regression coefficients express the expected change in the ratio $\gamma/(1-\gamma)$ , for a one standard deviation increase of the concerned predictor, holding all the other predictors fixed at a constant value. <sup>b</sup>Odds ratio 95% confidence interval (CI), lower bound. <sup>c</sup>Odds ratio 95% CI, upper bound. <sup>d</sup>Population intercept.

### Landscape genomics

#### Population structure analysis

After pruning for MAF, LD, genotype and individual call rates, PSD counted 12,925 SNPs and 1,355 individuals, among which 743 from Uganda, 131 European taurine, 158 African taurine, 195 sanga from outside Uganda, and 128 zebu cattle.

Sanga and zebuine ancestries were the most represented in Uganda. Particularly, on average the sanga component constituted 76% (±13%) of the individual ancestries, whereas the zebuine counted 18% (±13%), with more than half of the individuals showing a zebuine proportion >20%. Further, ~3% of African and European taurine genomic ancestry components was also observed. Genomic components showed spatial structure, the zebuine gene pool being more present in the North-East of the country, and the sanga in central and south-western Uganda (S10 Fig.) [22]. The African taurine ancestry component was detectable as background signal especially in the North-West and South-West, whereas European introgression was mostly observed in the South-West.

The first three principal components (PC_1_, PC_2_ and PC_3_, respectively) explained almost the totality of the variance within ADMIXTURE Q-scores for *K*=4; PC_1_ split the dataset between sanga and zebu gene pools, and PC_2_ and PC_3_ identified the European and African taurine components, respectively. Thus, these three PCs were used as population structure predictors to account for population structure within LGD in the landscape genomics models.

#### Genotype-environment associations

After QC, LGD counted 40,886 markers and 743 animals (the same in PSD) from 199 farms (4±1 samples/farm), over 51 cells (15±5 samples/cell).

Sixty-three genotypes across 41 putative adaptive loci resulted significantly associated with *Ψ_R_* (Fig 3A, S2 Table, and S11-S12 Figs). Eight genotypes across seven loci resulted significantly associated with *γ* (Fig 3B, S3 Table, and S11-S12 Figs).

**Fig 3.**
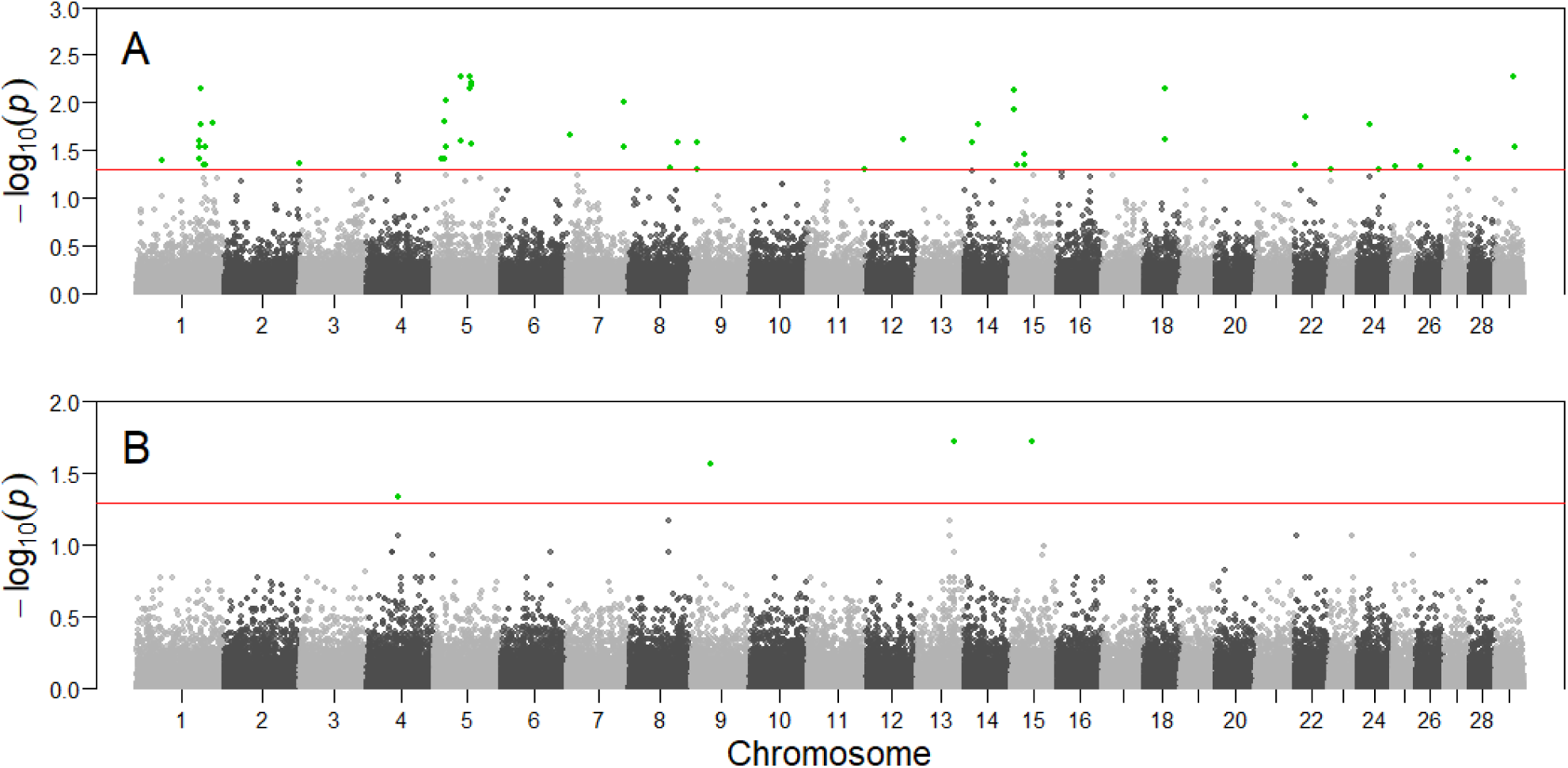
Manhattan plots of the genotype-environment associations. *X*-axis reports chromosomal position of the tested SNPs on cattle chromosomes. *Y*-axis reports the test statistic *p-values* (*p*) for the associations with *R. appendiculatus* occurrence probability (A), and with *T. parva parva* infection risk (B). *P-values* are displayed for each genotype after the Benjamini-Hochberg (BH) correction, and on the –log_10_ scale. Nominal significance threshold (*α*BH=0.05) is displayed as a red line, and significant *p-values* are represented in green.

#### Gene annotation

Among the 41 loci significantly associated with *Ψ_R_*, 18 presented at least one annotated gene in the Ensembl database in close proximity (Table 5A and S12 Fig.). Locus BTA-113604-no-rs (hereafter BTA-113604) is located ~12.5 kbp apart from the Protein kinase, cGMP-dependent, type I (*PRKG1*) gene on chromosome 26. *PRKG1* was already proposed as a candidate gene for tick resistance in South African Nguni cattle [72].

Six out of the seven loci significantly associated with *γ* presented at least one annotated gene within the selected window size (Table 5B and S12 Fig.). Two SNPs (ARS-BFGL-NGS-110102 and ARS-BFGL-NGS-24867, hereafter ARS-110102 and ARS-24867, respectively) were proximal to the Src-like-adaptor 2 (*SLA*2) gene on chromosome 13. *SLA*2 human orthologue encodes the Src-like-adaptor 2, a member of the SLAP protein family which regulates the T and B cell-mediated immune response [73]. Given *T. parva parva* known ability to promote the proliferation of T and B cells [74,75], we considered *SLA*2 as a second candidate gene for ECF local adaptation.

**Table 5.** Gene annotation for the loci significantly associated with *Ψ*_*R*_ (A) and *γ* (B). 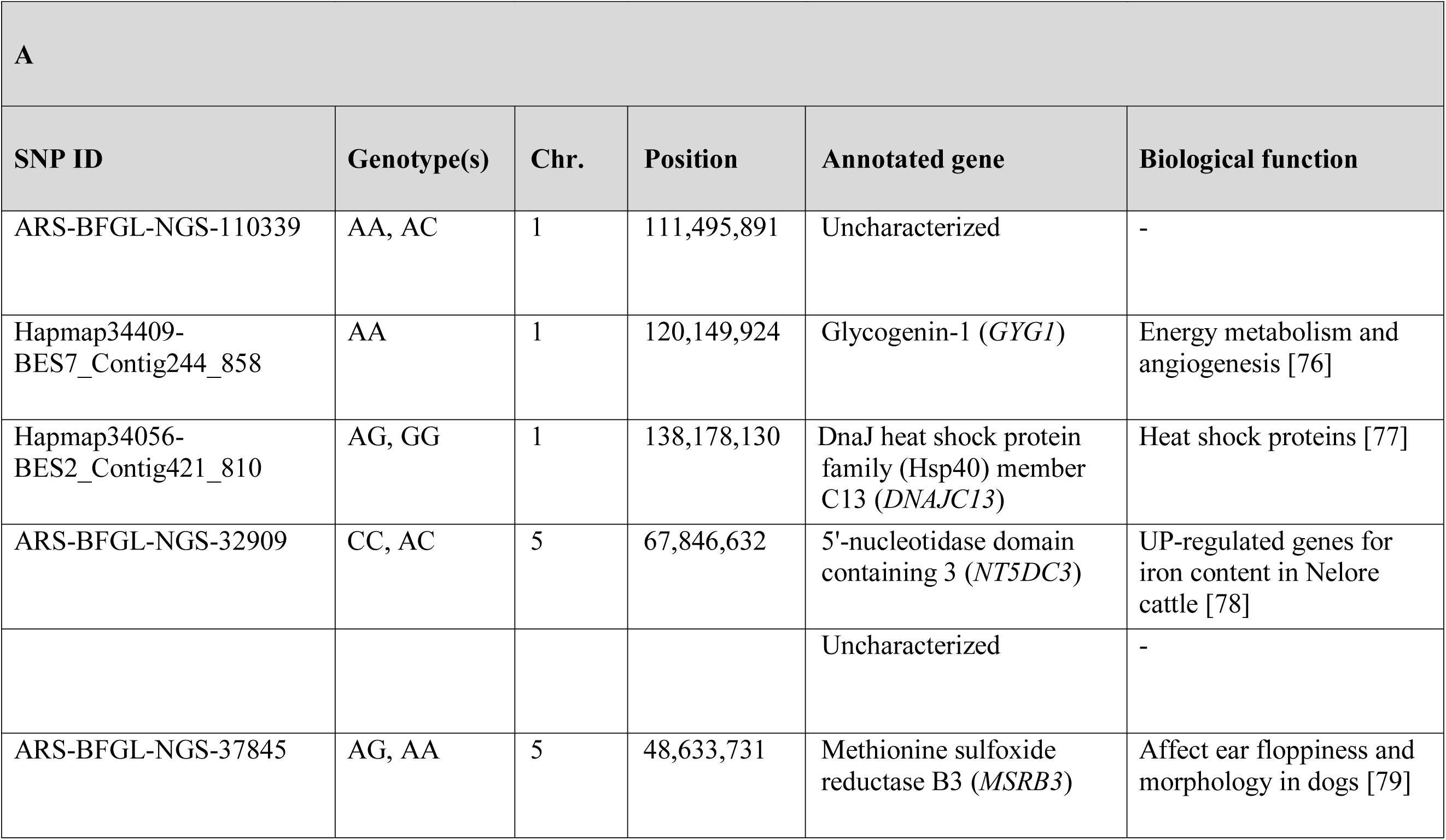

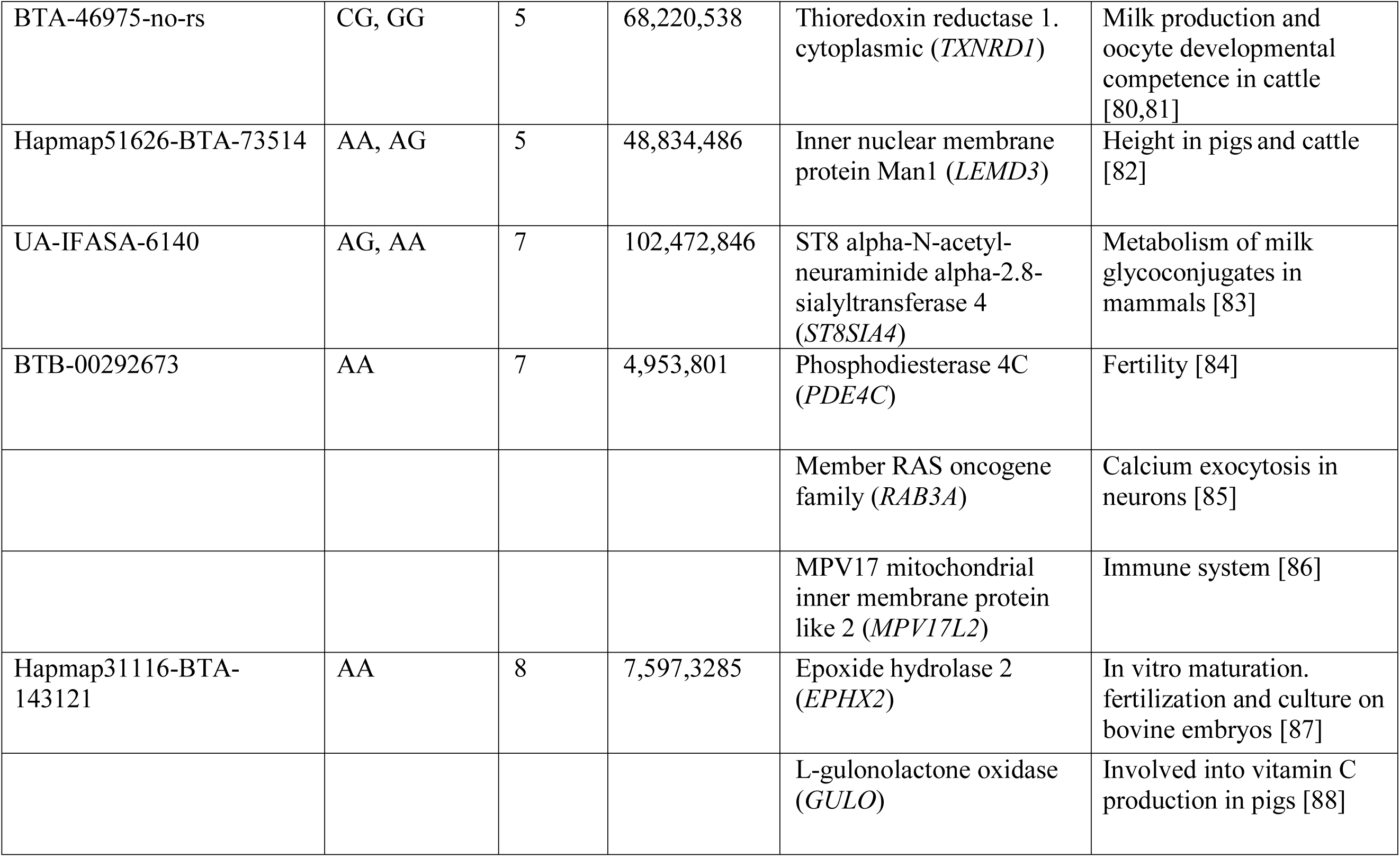

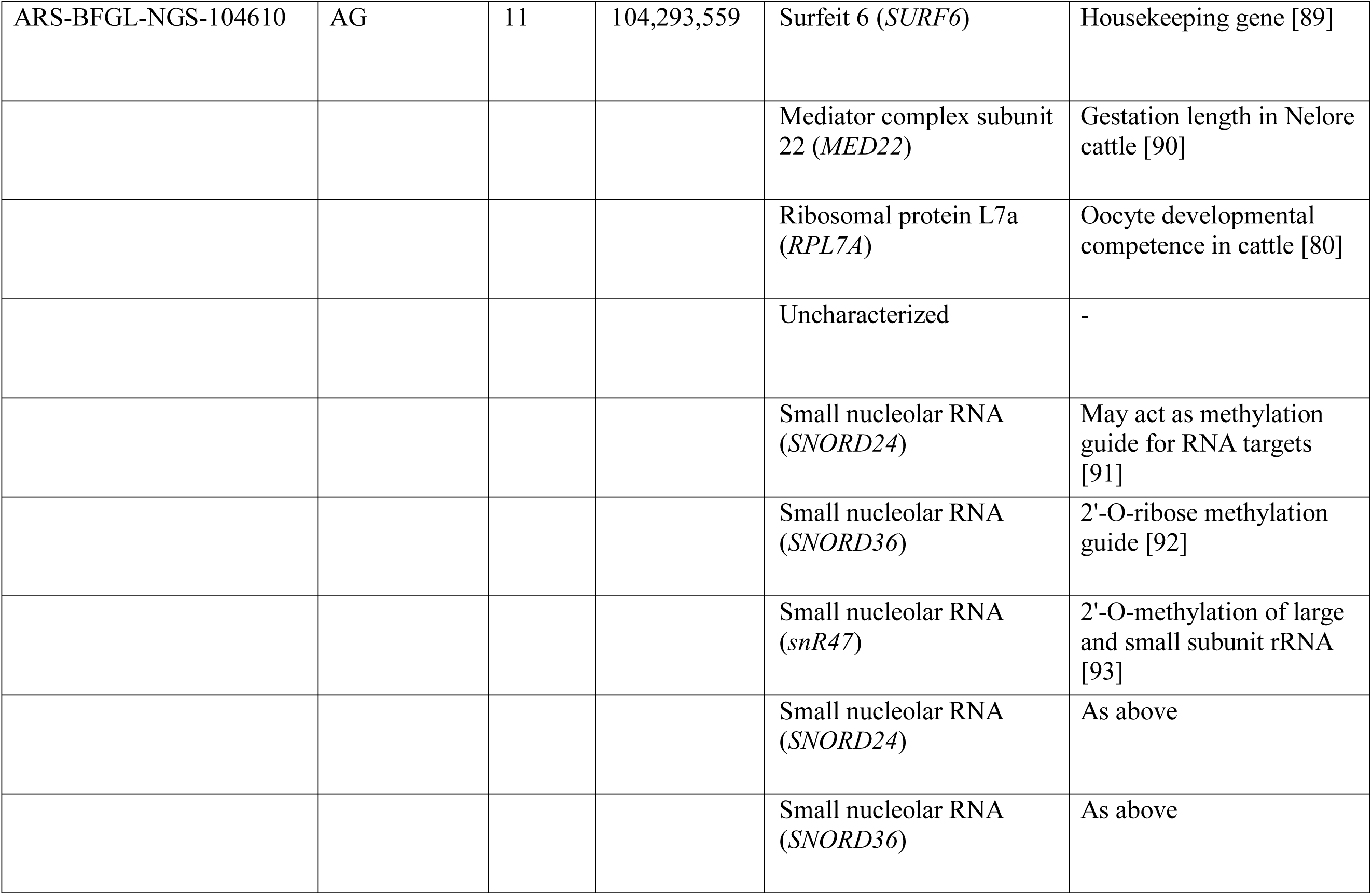

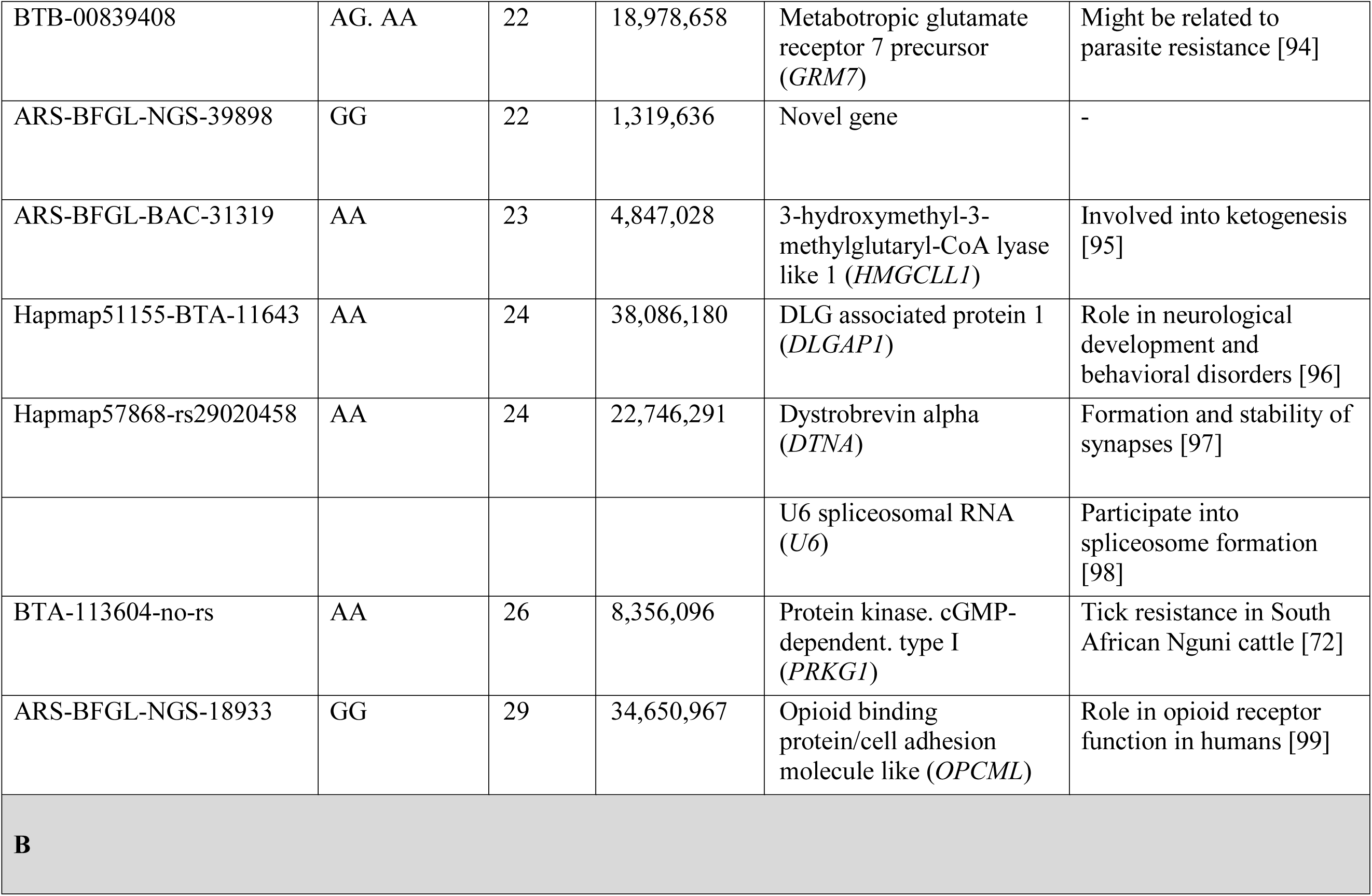

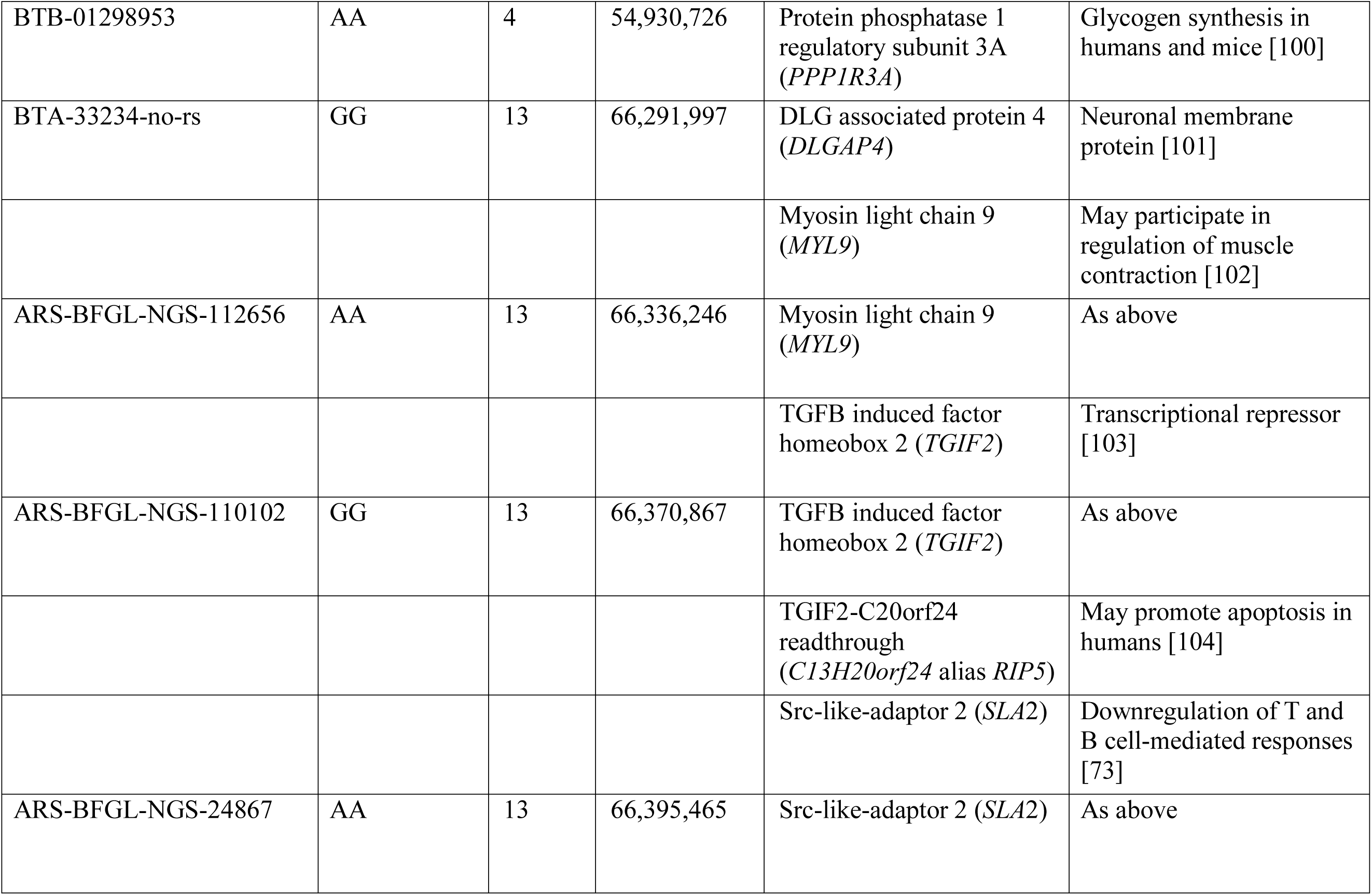

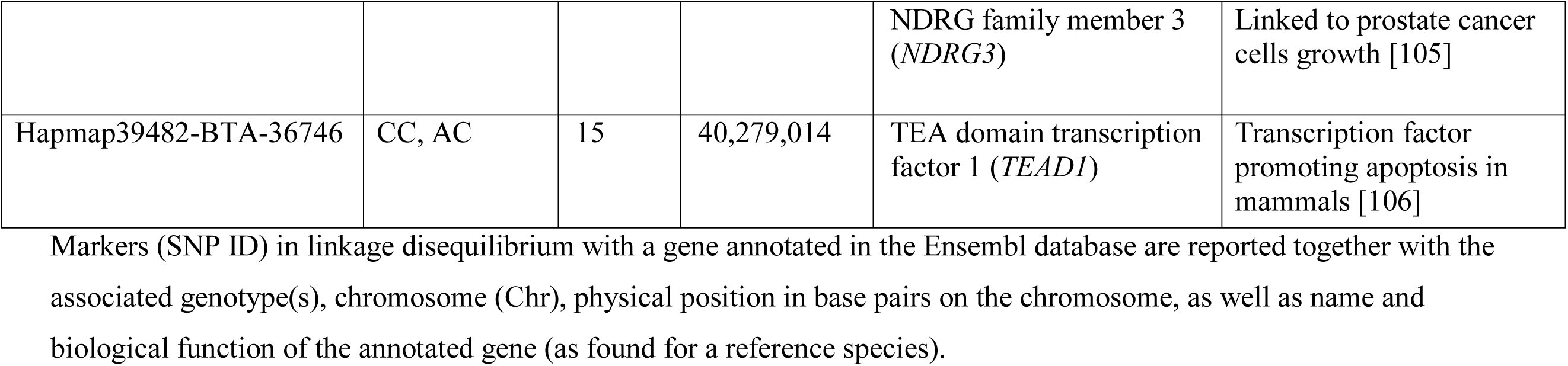

### Local ancestry

#### PCADMIX and Beta regression analyses

Based on the gene pools revealed by ADMIXTURE analysis in Ugandan indigenous cattle, we performed PCADMIX analysis using one zebuine (Tharparkar; THA) and one African taurine (Muturu; MUT) reference (S4 Text). After QC, LAD counted 689,339 markers and 128 individuals (102 Ugandan cattle individuals, 13 THA, and 13 MUT).

For the genomic window hosting BTA-113604 (i.e. window 13 on chromosome 26; S13 Fig.), 79 out of the 204 haploid individuals targeted showed MUT ancestry, while 125 THA ancestry (S14 Fig.). For the genomic window hosting ARS-110102 and ARS-24867 (i.e. window 145 on chromosome 13; S13 Fig.), 63 haploid individuals were assigned to MUT, while 141 to THA (S14 Fig.).

Tharparkar ancestry at window 13 of chromosome 26 showed a positive and significant association with *Ψ_Rc_* (*H_0_*: *β_i_*=0, *α*=0.05) (Table 6 and Fig 4), while no significant association was found between the Muturu/Tharparkar ancestries at window 145 of chromosome 13 and *γ_c_* (S5 Text).

**Fig 4.**
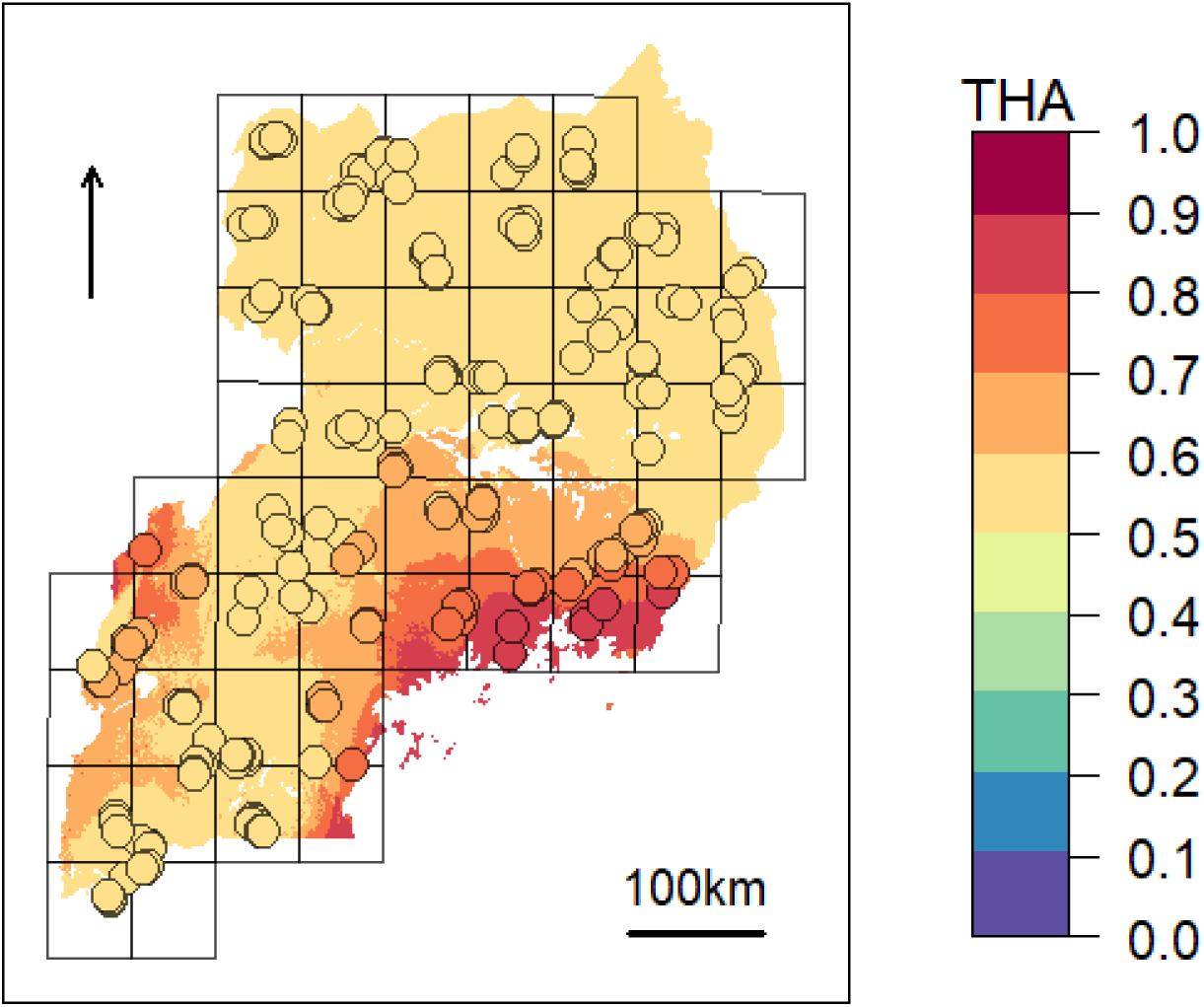
Expected zebuine proportion for the genomic region candidate for tick resistance over the study area. The association inferred through beta regression between Tharparkar ancestry (THA) and average *R. appendiculatus* occurrence probability per cell (Table 6) was used to generalize expected zebuine ancestry over the entire study area. Colour key corresponds to predicted THA proportion, with increasing values from blue to red tones. Sampled farms are represented with circles, and coloured according to the predicted THA proportion at their geographical location.

**Table 6.**
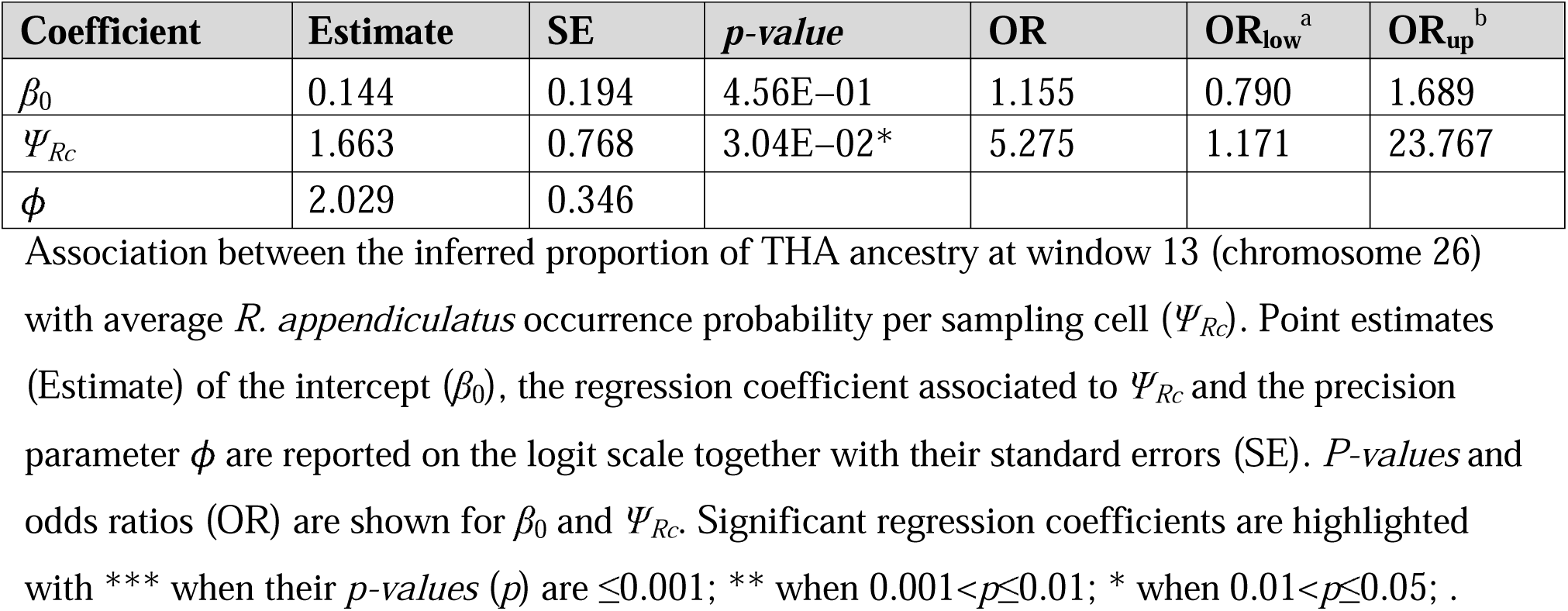
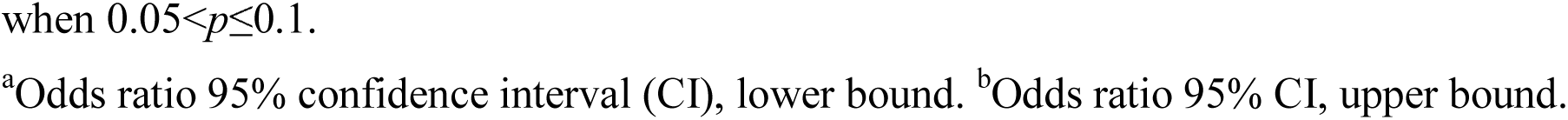
Beta regression results.

| Coefficient | Estimate | SE | *p-value* | OR | OR<sub>low</sub><sup>a</sup> | OR<sub>up</sub><sup>b</sup> |
| --- | --- | --- | --- | --- | --- | --- |
| $\beta_0$ | 0.144 | 0.194 | 4.56E-01 | 1.155 | 0.790 | 1.689 |
| $\Psi_{Rc}$ | 1.663 | 0.768 | 3.04E-02* | 5.275 | 1.171 | 23.767 |
| $\phi$ | 2.029 | 0.346 | | | | |

Association between the inferred proportion of THA ancestry at window 13 (chromosome 26) with average *R. appendiculatus* occurrence probability per sampling cell ( $\Psi_{Rc}$ ). Point estimates (Estimate) of the intercept ( $\beta_0$ ), the regression coefficient associated to $\Psi_{Rc}$ and the precision parameter $\phi$ are reported on the logit scale together with their standard errors (SE). *P-values* and odds ratios (OR) are shown for $\beta_0$ and $\Psi_{Rc}$ . Significant regression coefficients are highlighted with \*\*\* when their *p-values* (*p*) are $\leq 0.001$ ; \*\* when $0.001 < p \leq 0.01$ ; \* when $0.01 < p \leq 0.05$ ; .
when $0.05 < p \leq 0.1$ .
<sup>a</sup>Odds ratio 95% confidence interval (CI), lower bound. <sup>b</sup>Odds ratio 95% CI, upper bound.

## Discussion

East Coast fever represents a major issue for livestock health in sub-Saharan countries [107], with over one million cattle deceased every year, and an annual economic damage of 168-300 million USD [2,108].

ECF incidence is highly correlated with the geographical distribution of the tick vector *R. appendiculatus*, whose occurrence is an essential precondition for *T. parva parva* infection in cattle [3]. However, with the present study we show that areas with predicted poor habitat suitability for the tick can present higher infection rates when compared with regions highly suitable for the tick (Fig 2 and Table 4). Such observation suggests that *T. parva parva* occurrence cannot be explained by the sole presence of its vector. Here, we suggest three possible hypotheses to explain such a counterintuitive pattern.

First, environmental temperature may play a pivotal role in defining *T. parva parva* infection risk in cattle. Piroplasm development within the tick vector appears to be hindered by temperatures >28°C persisting even for short time periods (as less as 15 days) [54]. Therefore, areas exceeding this temperature threshold might present a reduced infection risk due to the low success in parasite development and transmission. The presence of such a temperature constraint might concur in explaining the low infection risk predicted in the regions such as North-East of Lake Victoria, where a highly suitable habitat is predicted for *R. appendiculatus*, but where temperature can reach 30°C in the warmest month of the year (January) [27]. Coherently, in the south-western area, environmental temperature ranges between ~8-28°C during the whole year [27], and the predicted risk of infection increases despite the decrease in habitat suitability for the tick.

Second, the most suitable areas for the vector overlap those regions where the highest levels of zebuine ancestry were recorded (S10 Fig.). *B. indicus* is known to be more effective in counteracting tick infestation than *B. taurus* [109–112], and is consequently less affected by tick-borne micro-organisms [111], including *T. parva parva*, whose effects are known to be dose-dependent [107,113]. The core adaptive response to tick burden was identified as the inflammatory reaction triggered by the tick bite at the cutaneous level [111], which activates a strong white cells-mediated cutaneous reaction [114] affecting attachment, salivation, engorgement, and ultimately limiting the inoculation of tick-borne microorganisms [115]. Therefore, the low infection risk observed in the most suitable areas for *R. appendiculatus* (e.g. north-eastern districts) might be explained by the coexistence of putative tick-resistant zebuine-like populations [116], along with a sub-optimal environmental niche for the parasite. Further, we speculate that cattle populations living in regions suitable for *T. parva parva* development, but with reduced predicted tick presence (e.g. the southern districts), could have not underwent a tick-specific adaptation, and therefore show higher infection rates.

Third, the *R. appendiculatus* distribution model does not explicitly consider anthropogenic factors like tick-control campaigns on a local and temporal basis. However, adequate tick-control campaigns are rarely undertaken in Uganda (Ugandan National Drug Authority), and evidence of *R. appendiculatus* developing drug resistance has been recorded [117].

Despite infections being observed in the northern farms of Uganda, an almost null occurrence of the ECF-vector is predicted for the same regions (0<*Ψ_R_*<0.1; Fig 2A). A possible explanation is the lack of *R. appendiculatus* records, and the consequent bias in the tick distribution model [37,118].

Moreover, predicted infection risk in the North (0<*γ*<0.3; Fig 2B) may be inflated by the global inverse relationship between *γ* and *Ψ_R_* as estimated by the infection risk model (Table 4), and care is recommended regarding the infection risk predictions for these areas.

Here, we suggest that the putative adaptive component sustaining ECF-endemic stability might be due to a synergic mechanism involving specific adaptations to *R. appendiculatus* (the vector) and *T. parva parva* (the parasite). Specifically, adaptations to tick burden could be found along the Lake Victoria coasts, where a higher selective pressure linked to *R. appendiculatus* occurrence is predicted (Fig 2A). We identified 41 loci across 18 chromosomes significantly associated with *Ψ_R_* (Fig 3A), with the majority of putative loci under selection found on the chromosomes 5 (nine loci), 1 (seven loci), and 15 (three loci). Interestingly, the large genomic region hosting the associated SNPs on chromosome 5 (S2 Table) overlaps a genomic region which has been previously associated with several traits in tropical cattle, including parasite resistance [119]. Among the genes in LD with the associated markers, we found *PRKG1* on chromosome 26 (Table 5A and S13 Fig.), a gene coding for an important mediator of vasodilation, and already reported as possibly involved in tick resistance in the South African Nguni breed (see Table 6 in [72]). Importantly, vasodilatation is a classical feature of the inflammatory response [120, 121], the core mechanisms underlying tick resistance, as discussed before. None of the remaining annotated genes was easily attributable to adaptation to tick burden (Table 5A).

A specific adaptive response towards *T. parva parva* infection may have evolved in south-western Uganda, possibly due to ecological conditions suitable for the parasite survival, and to the presence of a more tick-susceptible cattle population (S10 Fig.). *Theileria parva parva* pathogenicity is linked to its ability to invade host lymphocytes, and promoting their transformation and clonal expansion through the activation of several host-cell signalling pathways [15,75,122]. Here, we found seven markers significantly associated with *γ*, two of which (ARS-110102 and ARS-24867) included within *SLA*2 genic region on chromosome 13 (S13 Fig.). *SLA*2 is known to be involved with signal transduction in B and T cells. Further, *SLA*2 downregulates humoral and cell-mediated immune responses, by contributing to a correct lymphocytes’ activation and proliferation [73,123,124]. *SLA*2 antagonistic effect on lymphocytes proliferation would suggest its putative involvement in opposing the diffusion of *T. parva parva* in the organism. However, further molecular and immunological investigation will be required for validating this hypothesis.

Despite the genetic proximity between Muturu and some tick resistant indigenous *B. taurus* breeds of western Africa (i.e. N’Dama) [111,125], local ancestry of the genomic region candidate for tick resistance was predominantly assigned to Tharparkar (Fig 4, Table 6, and S14 Fig.). This result is in agreement with the known resistance of zebuine cattle to ticks, and suggests the origin of tick resistance in eastern Africa either from imported Indian populations or within local zebuine-like populations after migration from India. Conversely, no easily-interpretable indication was obtained for the genomic region candidate for tolerance to *T. parva parva* infection. Indeed, neither Tharparkar nor Muturu ancestries displayed a significant association with infection risk, while an additional local ancestry analysis revealed a positive correlation with the European taurine Hereford ancestry when tested versus Tharparkar (S5 Text). Although surprising, this result would rather point towards a taurine origin of infection tolerance. However, local ancestry results are inherently reference-dependent [66], and further analyses with different African taurine and zebuine references will be required to disentangle the evolutionary origin of the genomic regions under scrutiny.

Besides the identification of candidate regions for ECF local adaptation, our results revealed allochthonous introgression from Europe within the local gene pools of Ugandan cattle (S4 Text and S10 Fig.). This finding is consistent with the generalized loss of agro-biodiversity reported worldwide [8,126], and stresses the importance of monitoring local genetic resources to conserve unique adaptations, including tolerance to tropical endemic diseases.

Despite limitations in both epidemiological and species occurrence data, the proposed models allowed the identification of two candidate genes for ECF-tolerance. In general, the combination of ecological modelling (e.g. species distribution models) and landscape genomics showed the potential of revealing candidate genomic regions for local adaptation, and could be considered in any evolutionary study involving interacting species, like symbiotic relationships (i.e. mutualism, parasitism and commensalism), and competition.

## Acknowledgments

The authors are grateful to the members of the NextGen Consortium (http://nextgen.epfl.ch), and to Graeme S. Cumming, who kindly provided the *R. appendiculatus* occurrence dataset used in the present study.

## Data Accessibility

Raster data are available from the public sources mentioned in the references and in S1 Text. Datasets and source code are available from the Dryad Digital Repository (doi:10.5061/dryad.sf5j2bf).

## Author Contributions

Conceptualization: EV, MB, LC, MM, ER, EF, MDC, RN, SJ, PAM. Data Curation: EV, MB, LC, MM. Formal Analysis: EV, MB. Methodology: EV, MB, ER, EF, SJ, PAM. Software: EV, MB, MM, MDC. Visualization: EV. Original Draft Preparation: EV. Review and Editing: MB, LC, MM, ER, SJ, TSS, PAM. Investigation: LC, MM, FK, RN. Funding Acquisition: LC, RN, SJ, PAM. Supervision: LC, SJ, PAM. Project Administration: SJ, PAM. Resources: CMK, CMS, VM, FK, TSS, HJH.

## Supporting information

**S1 Text. MODIS collection 5 data.**

**S2 Text. Statistical significance of genotype-environment associations.**

**S3 Text. Additional SAMβADA analysis (*K*=3 and *K*=16 corrections).**

**S4 Text. Selection of reference populations for local ancestry analyses.**

**S5 Text. Beta regression results on additional local ancestry analyses.**

**S1 Fig. Bioclimatic variables used in *R. appendiculatus* distribution model.**

**S2 Fig. Selection of the annual period with NDVI values best explaining *S. caffer* records.**

**S3 Fig. Outlier detection in the infection model predictors.**

**S4 Fig. Additional ADMIXTURE analysis: results.**

**S5 Fig. Correlations between bioclimatic variables and principal components.**

**S6 Fig. Candidate *R. appendiculatus* distribution models and model selection.**

**S7 Fig. Selected *R. appendiculatus* distribution model with confidence intervals.**

**S8 Fig. Candidate *S. caffer* distribution models and model selection.**

**S9 Fig. Selected *S. caffer* distribution model with confidence intervals.**

**S10 Fig. Map of the Ugandan ancestry components as derived by ADMIXTURE analysis (*K*=4).**

**S11 Fig. Quantile-Quantile plots of the genotype-environment association studies (*K*=4 correction).**

**S12 Fig. Additional SAMβADA analysis (*K*=3 and *K*=16 corrections): results.**

**S13 Fig. *PRKG1* and *SLA2* genomic regions.**

**S14 Fig. Spatial representation of PCADMIX assignments (Tharparkar/Muturu comparison).**

**S1 Table. Composition of the population structure dataset.**

**S2 Table. SAMβADA analysis (*K*=4 correction): significant associations with *R. appendiculatus* occurrence probability.**

**S3 Table. SAMβADA analysis (*K*=4 correction): significant associations with *T. parva parva* infection risk.**

